# Pathological R-loops in bacteria from engineered expression of endogenous antisense RNAs whose synthesis is ordinarily terminated by Rho

**DOI:** 10.1101/2024.04.20.590381

**Authors:** Apuratha Pandiyan, Jillella Mallikarjun, Himanshi Maheshwari, Jayaraman Gowrishankar

## Abstract

In many bacteria, the essential factors Rho and NusG mediate termination of synthesis of nascent transcripts (including antisense RNAs) which are not being simultaneously translated. It has been proposed that in Rho’s absence toxic RNA-DNA hybrids (R-loops) may be generated from nascent untranslated transcripts; and genome-wide mapping studies in *Escherichia coli* have identified putative loci of R-loop formation from more than 100 endogenous antisense transcripts that are synthesized only in a Rho-deficient strain. Here we provide evidence that engineered expression in wild-type *E. coli* of several such individual antisense regions on a plasmid or the chromosome generates R-loops that, in an RNase H-modulated manner, serve to disrupt genome integrity. Rho inhibition was associated with increased prevalence of antisense R-loops also in *Xanthomonas oryzae* pv. *oryzae* and *Caulobacter crescentus*. Our results confirm the essential role of Rho in several bacterial genera for prevention of toxic R-loops from pervasive yet cryptic endogenous antisense transcripts. Engineered antisense R-looped regions may be useful for studies on both site-specific impediments to bacterial chromosomal replication and the mechanisms of their resolution.

## Introduction

The Rho protein mediates an important and active mechanism of bacterial transcription termination, and is essential for viability in many species including *Escherichia coli* [reviewed in (1–7)]. Rho binds to an exposed *r*ho *ut*ilization (*rut*) sequence of degenerate specificity on nascent RNA trailing an RNA polymerase (RNAP) during the elongation step of transcription, following which it acts along with accessory factors such as NusG to dislodge RNAP from the DNA; the precise molecular mechanism by which Rho’s action leads to transcription termination and to release of RNAP and RNA from DNA is unclear, and several alternative models have been proposed in this regard (8–14).

In view of the existence of transcription-translation coupling in bacteria such as *E.coli* (1, 15–19) for which NusG is also proposed to play a role (20, 21), *rut* sequences in the protein-coding regions of mRNAs are shielded from Rho binding, so that Rho mediates premature termination of only those mRNAs that are not being simultaneously translated (such as those with a nonsense codon mutation in a protein-coding gene or on which there has been stochastic failure of ribosome loading); such premature termination within a gene or operon as a consequence of transcription-translation uncoupling is also referred to as transcriptional polarity [(22–24); reviewed in (1, 15, 16, 22, 25)]. Rho also mediates transcription termination of RNAs that are not naturally translated (such as in the 3’-untranslated region of an mRNA, or antisense RNAs), although an active antitermination mechanism operates to ensure that synthesis of ribosomal RNA (which is not translated) is not terminated by Rho’s action.

In *Bacillus subtilis*, transcription and translation are not strongly coupled (26, 27), and it has been proposed that *cis*-encoded elements define the sites of Rho-mediated transcription termination (26, 28). At the same time, the finding from *E. coli* (29, 30) that Rho acts to suppress pervasive antisense transcription has been shown also to be true in the different Gram-positive bacteria including *B. subtilis* (26, 31), *Staphylococcus aureus* (32), and *Mycobacterium tuberculosis* (33, 34). For *E. coli*, several models, not mutually exclusive, have been proposed to account for the essentiality of Rho, such as its role in (a) silencing of cryptic prophage genes (35–38), (b) prevention of replication fork arrest especially at sites of stalled transcription elongation complexes (39–41), or (c) avoidance of formation of toxic transcription-associated RNA-DNA hybrids or R-loops in the genome (25, 29, 42–44).

The R-loop is a three-stranded nucleic acid structure that is generated when a nascent transcript is re-annealed with the upstream template DNA strand to form an RNA-DNA hybrid, with the non-template strand now being displaced as single-stranded (ss)-DNA [reviewed in (44, 45)]). RNase HI (encoded by *rnhA*) specifically cleaves RNA in an RNA-DNA hybrid. R-loops can confer cellular toxicity by one or more of the following mechanisms: (a) by acting as roadblocks to transcription (29, 46, 47); (b) by engendering transcription-replication conflicts (48–51); and (c) by priming aberrant DNA replication initiation (52–57).

Two mechanisms, not mutually exclusive, may be envisaged for R-loop generation from nascent untranslated RNAs in the absence of Rho-mediated transcription termination. First, a transcript may by itself directly re-anneal with the template DNA strand in the negatively supercoiled region upstream of the moving RNAP (25, 42, 44). Second, RNAP in a transcription elongation complex that is uncoupled from translation is more prone to backtracking and arrest (40, 58, 59), and there is evidence for R-loop formation from its associated RNA when the elongation complex collides with a replisome (40, 48, 49, 60–62).

Support for the R-loop avoidance model of Rho and NusG essentiality in *E. coli* has emerged from the findings (a) that nucleic acid preparations from cells with compromised Rho function exhibit increased reactivity to S9.6 monoclonal antibody, which is specific for RNA-DNA hybrids (29); (b) that Δ*rho* and Δ*nusG* lethalities are rescued by ectopic expression of an R-loop helicase UvsW (sourced from phage T4) (29, 43); and (c) that many novel antisense RNAs are exclusively detected in a Δ*rho* strain which has been rendered viable with UvsW helicase, and whose coding locations are correlated with the hotspots of antisense R-loop formation that have been mapped in the genome (29). These data have been interpreted to suggest that several hundred cryptic antisense loci exist in the *E. coli* genome whose expression is ordinarily terminated by Rho and which, in absence of Rho, serve as sites for formation of toxic R-loops that are correlated with Δ*rho* lethality. Several other studies in *E. coli* (63–66) and *Vibrio cholerae* (67) have also suggested that R-loops are generated when transcription and translation are uncoupled. However, these lines of evidence are not conclusive, since UvsW belongs to a group of helicases that can act not only on R-loops but also at replication forks (68–71), and furthermore Rho deficiency itself can confer pleiotropic effects on cells (2).

In the present work, we show that the engineered overexpression in *E. coli* of several of these otherwise cryptic endogenous antisense loci on plasmids or the chromosome confers toxicity and loss of genome integrity even in Rho-proficient strains; the effects are RNase H-modulated, indicating that R-loops are being generated under these conditions (72). Our findings therefore confirm the existence of significant numbers of R-loop forming natural antisense RNAs whose synthesis is being ordinarily curtailed by Rho. We have also extended these findings to two other Gram-negative bacteria, *Xanthomonas oryzae* pv. *oryzae* and *Caulobacter crescentus*, to show in them that Rho inhibition is associated with increased R-loop prevalence that can putatively be mapped to antisense hotspots across the genome. Finally, our work offers a new experimental approach based on transcription-associated R-loops to provoke site-specific impediments to replisome progression on the chromosome, which is different from those based on protein roadblocks (73, 74), ectopic *Ter* sites (75), palindrome DNA cleavage (76), and *rrn* operon inversions (60) or other head-on transcription (48) that have been described earlier.

## Materials and Methods

### Bacterial strains and growth conditions

*E. coli* strains are listed in Supplementary Table S1, and their derivatives were grown in LB medium (77) at 37° unless otherwise indicated. The knockout mutants Δ*rnhA*, Δ*pcnB*, Δ*entE*, Δ*cstA,* and Δ*ilvD* were from the Keio collection of BW25113 derivatives (78). Where appropriate, the medium was supplemented with ampicillin (Amp), kanamycin (Kan) or trimethoprim (Tp) (each at 50 μg/ml), and Xgal at 25 μg/ml. Supplementation with isopropyl β-D-thiogalactoside (IPTG) was at 1 mM.

*C. crescentus* and *X. oryzae* pv. *oryzae* strains were provided by Anjana Badrinarayanan and Subhadeep Chatterjee, respectively, and their growth conditions were as described (79, 80). Supplementation of cultures with the Rho inhibitor bicyclomycin (BCM) for each of the bacteria was at the indicated concentrations.

### Plasmids

Previously described plasmids include (salient features in parentheses): pKD13 (Kan^R^ Amp^R^), pKD46 (Amp^R^), and pCP20 (Cm^R^ Amp^R^) for chromosomal recombineering and site-specific excision between a pair of FRT sites, as described (81); pTrc99a (Amp^R^, for IPTG-induced expression of cloned DNA fragments) (82); pSK760 (Amp^R^, ColE1 plasmid with cloned *rnhA^+^* gene) (63); pET28b (Kan^R^, ColE1-derived plasmid) (Novagen, USA); and pHYD5701 (Tp^R^, single-copy-number plasmid with *recA^+^ lacZ^+^*) (83).

Plasmids pHYD7527 and pHYD7528 are pKD13 derivatives with a cloned P*_tac_* sequence at a single site in two orientations, whose construction is described below. Construction of plasmids by PCR and cloning of different antisense regions into the pTrc99a vector is described in the Supplementary Text; the plasmids are listed in Supplementary Table S2, and the oligonucleotide primers used for generating PCR amplicons for the cloning are listed in Supplementary Table S3.

### Plasmid DNA supercoiling assays

Transformants of XL1-Blue (84) with pTrc99a-derived plasmids were grown to an *A*_600_ of around 0.6 without or with terminal exposure to IPTG for 2 h. Approximately 2 μg of plasmid DNA from each culture was subjected to electrophoresis at 3V/cm for 16 to 22 h on 1% agarose gels with the indicated concentrations of chloroquine, as described (55, 85).

### Recombineering of P*_tac_* at different chromosomal loci

We first cloned the P*_tac_* sequence in two orientations into plasmid pKD13 by annealing a pair of complementary oligonucleotide primers JGAH068-JGAH069 comprising the *tac* sequence (Supp. Table S3) and ligating the DNA duplex (with 5’-GATC overhangs) into the *BamHI* site of pKD13. From the transformants recovered in strain S17-1 1*_pir_* (86), two plasmids pHYD7527 and pHYD7528 were identified to carry the cloned P*_tac_* sequence reading, respectively, away from or towards the Kan_R_ gene of pKD13. The orientation of P*_tac_* in the plasmids was established by PCR with primer pairs JGAH070-JGAH072 (positive for pHYD7527) and JGAH071-JGAH072 (positive for pHYD7528); the primer sequences are given in Supplementary Table S3.

Plasmids pHYD7527 and pHYD7528 were then used as templates for recombineering of P*_tac_* into different sites on the chromosome, the details of which are described in the Supplementary Text. Expression of the λ Red genes for recombineering was achieved with the use of either plasmid pKD46 (81) or a strain carrying mini-λ-Tet prophage (87).

### Blue-white assay to determine RecA-dependence for viability of recombineered strains

The blue-white assay, that was employed to establish whether the P*_tac_*-recombineered strains were RecA-dependent or –independent for viability, has been described earlier (83, 86). Briefly, P*_tac_*-recombineered derivatives of XL1-Blue (which is *lacZ*ΔM15 and *recA1*) carrying the single-copy-number *recA^+^ lacZ^+^* Tp^R^ shelter plasmid pHYD5701 were plated for single colonies on LB-Xgal medium. Since the plasmid is spontaneously lost from approximately 20% of cells in a culture, RecA-independence of viability was established by the presence of white colonies on plating. On the other hand, if blue colonies alone were obtained (despite absence of Tp in the plates), it was concluded that the recombineered strain is RecA-dependent for viability.

### Whole genome sequencing (WGS) for DNA copy number analysis and mutation identification

The methods for WGS and analysis were essentially as described (83, 86). Genomic DNA was extracted by the phenol-chloroform method from cultures grown with IPTG supplementation to early-exponential phase, and subjected to Illumina-based WGS (Medgenome Labs, Bengaluru, India) followed by copy number analysis after alignment to the MG1655 reference sequence (Accession number U00096.3). WGS data were also used for de novo contig assembly and identification of novel junction sequences as described in Durrant et al (88).

### WGS following bisulphite treatment

The protocols for preparation of total nucleic acids from cultures, treatment (without denaturation) with sodium bisulphite, and subsequent whole-genome amplification, were as described (43). (Exposure to bisulphite provokes C-to-T changes in ss-DNA.) WGS was performed on an Illumina platform, and the reads were aligned and quantitated (using individual genes or ORFs as units) with the aid of the Bismark software (89); the reference sequences used were those of *E. coli* strain MG1655, *X. oryzae* pv. *oryzae* strain BXO1, and *C. crescentus* CB15 (Accession numbers U00096.3, NZ_CP033201.1, and AE005673.1, respectively). The Bismark algorithm determines, from the aggregate population of reads, base counts for each C residue of the reference sequence in the following categories: (i) unchanged, (ii) converted to T on top strand (“C-to-T”), or (iii) converted to T on bottom strand (“G-to-A”). For each ORF, the bisulphite reactivity of its top and bottom strands was then computed as the ratio of its C-to-T and G-to-A base counts, respectively, to its total base counts (that is, the sum of unchanged, C-to-T and G-to-A base counts).

### Other methods

Procedures for phage P1 transduction (90), pCP20-mediated excision between FRT sites (81), immunoblotting of total nucleic acid preparations with S9.6 monoclonal antibody (29), and determinations of β-galactosidase specific activity in Miller units (77) were as described. Methods for DNA manipulations, PCR and gel electrophoresis were as described in Sambrook and Russell (91).

## Results

### R-loop toxicity following plasmid-borne expression of endogenous chromosomal cryptic antisense regions

Previous studies have shown that R-loops can be generated from endogenous *E. coli* transcripts when expressed from plasmids in topoisomerase I-deficient cells but not in a wild-type strain [reviewed in (45)]. In wild-type *E. coli*, R-loops occur upon heterologous expression of certain mammalian gene regions such as from either an immunoglobulin or a triplet-repeat locus (both of which are known to form R-loops in their native contexts) (92, 93). Data from the experiments below indicate that plasmid-borne expression of several endogenous cryptic antisense regions can also generate R-loops in wild-type *E. coli*.

Twenty-two chromosomal antisense regions of *E. coli* (each approximately 0.3 kb or 2 kb long) were cloned downstream of the IPTG-inducible *trc* promoter of plasmid pTrc99a (82), as described in Supplementary Table S2. The regions were chosen from amongst those above the 85th percentile for antisense bisulphite reactivity (that is, for occurrence of displaced ss-DNA) (43). As discussed below, this criterion merely represents a first approximation for sites of antisense transcription leading to R-loop formation, since the percentiles have been determined across discrete ORFs whereas antisense transcription may be confined to only one part of an ORF or may even traverse ORF boundaries. Three additional antisense regions, that were each also expressed only upon Rho inhibition but were associated with a low bisulphite reactivity percentile and were therefore predicted not to exhibit R-loop propensity, were also chosen as controls in the experiments (Supp. Table S2). In the descriptions below, individual antisense regions are designated by the “b” number of the corresponding gene with “AS_” as prefix (94).

That a particular phenotype is R-loop associated can be inferred based on its modulation by cellular RNase HI status, that is, between derivatives that are Δ*rnhA*, *rnhA^+^*, or with engineered increase in RNase HI expression (40, 42, 48, 50, 63–65, 67, 92, 93). Of the 22 antisense regions that were tested in *recA^+^* derivatives of strain XL1-Blue after cloning in pTrc99a, six (AS_b0059, AS_b0397, AS_b0797, AS_b1077, AS_b2456, and AS_b2723) conferred a marked IPTG-dependent decrease in viability on Amp-supplemented medium, but only so in a Δ*rnhA* derivative and not the *rnhA^+^* parent (Fig. 1A, clone numbers marked in blue; only two of the clones that were negative in this test are shown, marked in black). We interpret these findings as evidence for R-loops being generated from the cognate antisense transcripts, whose persistence in the RNase HI-deficient strain then leads to inhibition of plasmid replication. None of the three control regions conferred growth inhibition in the *rnhA^+^* or Δ*rnhA* derivatives on IPTG-supplemented medium (data for two of them, AS_b0561 and AS_b2375, are shown in Fig. 1A, clone numbers marked in green).

**Figure 1:**
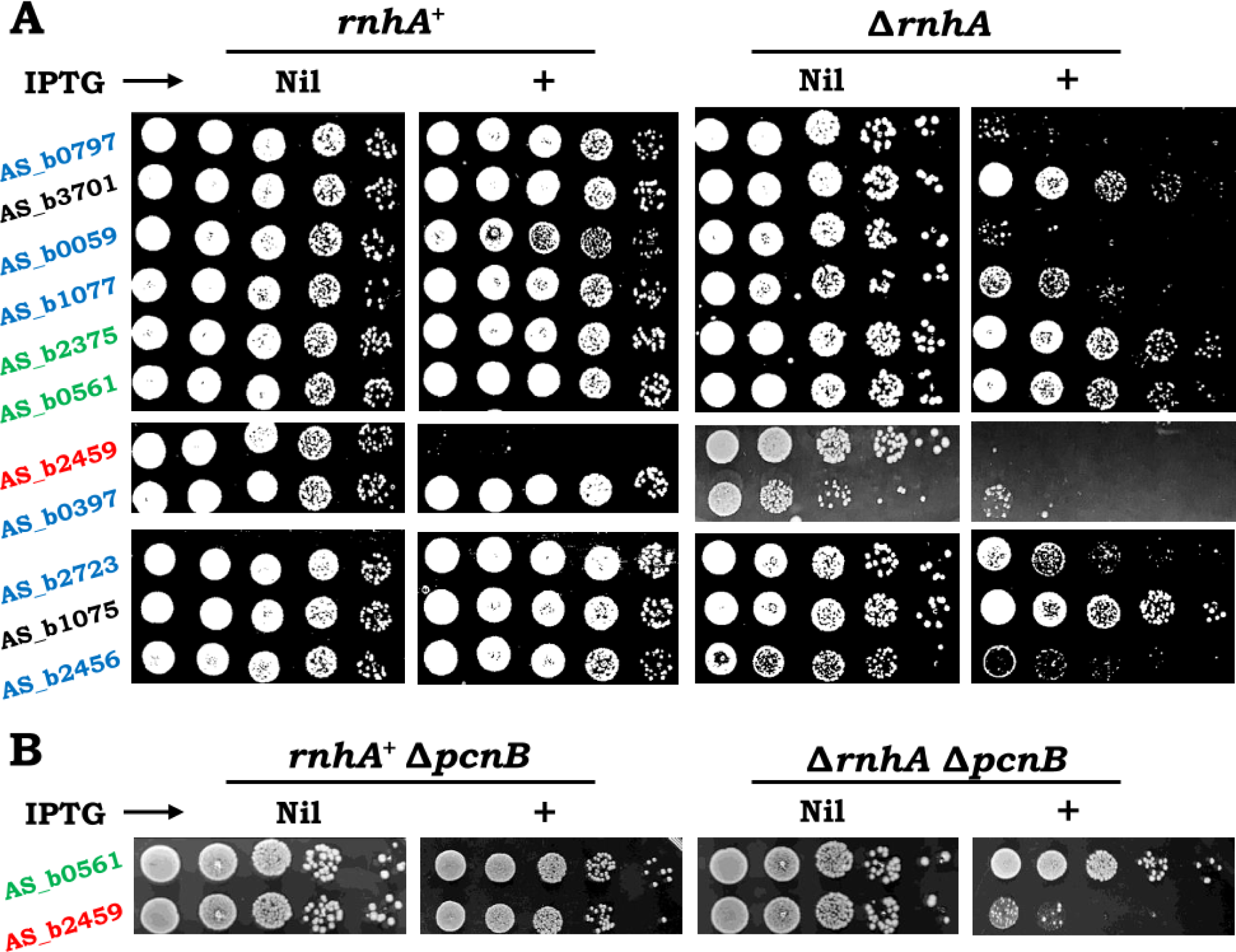
(**A**, **B**) Dilution-spotting assays of derivatives of XL1-Blue/pHYD5701 (*recA^+^*) with other alleles (*rnhA*, *pcnB*) as indicated, after transformation with pTrc99a derivatives carrying different cloned *E. coli* antisense regions. Spotting was done on Amp-supplemented plates without or with IPTG addition, as marked. Antisense region carried on each plasmid is given at left, with the following colour codes for different phenotypes (in *pcnB^+^* background): green, control; black, viable in all conditions tested; blue, IPTG-sensitive in Δ*rnhA*; and red, IPTG-sensitive in both *rnhA^+^* and Δ*rnhA*. AS_b0059 refers to AS_b0059 (region 1) in Supp. Table S2. Strains employed were pHYD5701-bearing derivatives of: *rnhA^+^*, XL1-Blue; Δ*rnhA*, GJ20007; ΔpcnB *rnhA^+^*, GJ20001; and Δ*pcnB* Δ*rnhA*, GJ20009.

For one more of the candidate antisense regions cloned in pTrc99a (AS_b2459, cloned region length 336 bp), IPTG-induced lethality on Amp-supplemented medium was observed in both Δ*rnhA* and *rnhA^+^* host strains (Fig. 1A, clone number marked in red). We surmise that in this case, R-loops generated from the antisense RNAs on the plasmids are in sufficient excess to overwhelm the capacity of native RNase HI to remove them. Even for this plasmid, modulation of IPTG-dependent lethality by cellular RNase HI status could be demonstrated following introduction into the host strain of a Δ*pcnB* mutation [which serves to reduce copy number of ColE1 plasmids (42, 95, 96)]; thus, the plasmid conferred lethality with IPTG in a Δ*rnhA* Δ*pcnB* but not an *rnhA^+^* Δ*pcnB* derivative (Fig. 1B, compare with control plasmid AS_b0561).

IPTG-dependent lethality associated with the plasmids above was exhibited also on medium without Amp supplementation (data not shown). These findings suggest that excessive R-loop formation on the plasmid not only can interfere with plasmid replication but also can confer cellular toxicity.

Notably, we found in several instances that genomic antisense regions conferring lethality upon transcription in pTrc99a are rather sharply defined, such that contiguous regions (< 1-2 kb away) are not lethal even though they too fulfil the candidate criteria for R-loop propensity. Three such examples, that reflect the approximations referred to above in choice of the antisense regions, were: AS_b1075 and AS_b1077; AS_b0059 (region 1) and AS_b0059 (region 2); and the group of AS_b2456, AS_b2458, and AS_b2459.

### IPTG-induced increased supercoiling in plasmid with cloned antisense region AS_b2459

Plasmid DNA supercoils are constrained upon R-loop occurrence, and the consequent homeostatic adjustment of superhelical density within the cells is manifested as increased negative supercoiling in preparations of plasmid DNA from these cultures that can be detected by chloroquine-agarose gel electrophoresis (45, 63–66, 72, 97–101). We tested the supercoiling status of the pTrc99a-derived plasmids prepared from cultures of XL1-Blue transformants grown without or with IPTG supplementation. Upon electrophoresis through a chloroquine-agarose gel, a marked increase in negative supercoiling following IPTG addition was observed for the plasmid with cloned antisense locus AS_b2459 (that confers lethality with IPTG supplementation even in an *rnhA^+^* strain), whereas supercoiling of the control plasmid with antisense region AS_b2375 was unaltered in absence or presence of IPTG (Fig. 2, compare lane 4 with lanes 1-3; gel image is shown in panel A and densitometric tracing of gel lanes in panel B). The lengths of cloned regions in the two plasmids are 336 bp and 334 bp, respectively. Similar results obtained from additional independent experiments are shown in Supplementary Figure S1. These findings serve to substantiate the conclusion that it is R-loop generation which is responsible for IPTG-induced lethality with AS_b2459.

**Figure 2:**
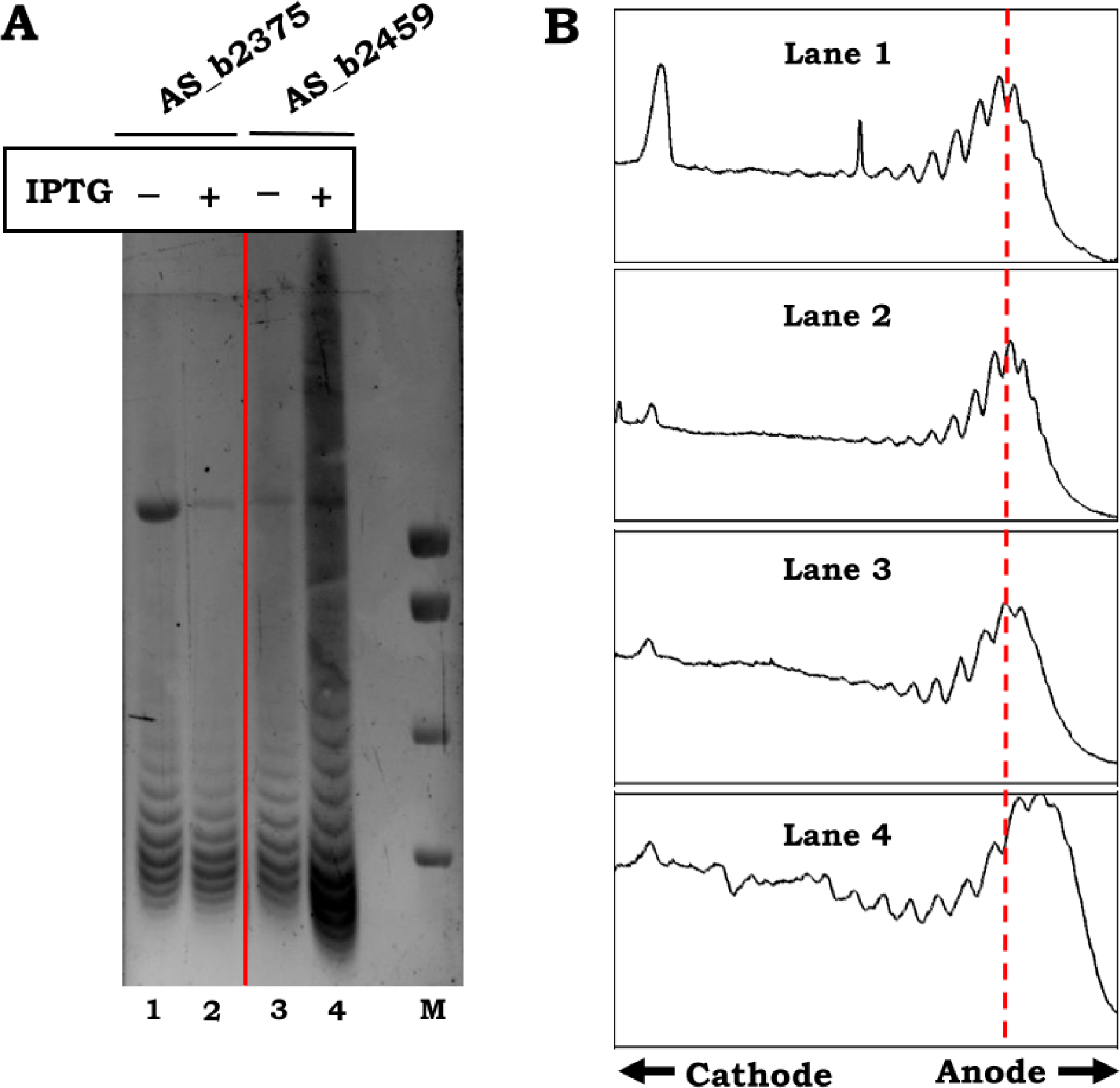
Supercoiling status of pTrc99a derivatives with cloned antisense regions AS_b2459 (test) and AS_b2375 (control) in cultures of XL1-Blue derivatives without and with IPTG supplementation. Gel electrophoresis was on 1% agarose with 4 μg/ml chloroquine; under these conditions, the more negatively supercoiled topoisomers migrate faster through the gel. (**A**) Gel image following ethidium bromide staining. M, lane for DNA size standards of 10, 8, 6, and 5 kb. (**B**) Densitometric traces of image along each of the indicated lanes. Vertical interrupted line is drawn to demonstrate that topoisomers with increased supercoiling are present in the population of AS_b2459-carrying plasmids prepared from culture with IPTG supplementation.

### R-loops from engineered antisense transcription at chromosomal locus AS_b2459

We chose two antisense regions AS_b2459 and AS_b0397, whose expression from the plasmid was lethal, respectively, even in *rnhA^+^* or only in Δ*rnhA*, to recombineer the P*_tac_* promoter sequence at their native chromosomal locations. The results for the two loci are described separately in this and the succeeding sections, respectively, but the following features were common to both of them.

All derivatives were Δ*lacZ recA1* on the chromosome and carried an unstable single-copy-number “shelter” plasmid with *recA_+_* and *lacZ^+^* genes; cultures of such derivatives typically are comprised of a 80:20 ratio (approximate) of plasmid-bearing and plasmid-free cells that grow as blue (*recA^+^*) and white (*recA1*) colonies, respectively, on Xgal-supplemented plates (“blue-white” assay).

Furthermore, although the P*_tac_* promoter used for the chromosomal recombineering was expected to be tightly IPTG-regulated, we observed in a control experiment (wherein this promoter was integrated upstream of native *lacZ* in each of two different constructs) that it is sufficiently active even in absence of IPTG (β-galactosidase specific activity of approximately 240 and 1980 Miller units, respectively, without and with 1 mM IPTG; details are provided in the Supplementary Text). The phenotypes described below were unchanged in presence or absence of IPTG.

For the antisense region AS_b2459, upstream chromosomal integration of the promoter was achieved by two different methods: in the first, the integration event results directly in the promoter reading outward into the antisense region, whereas in the second (two-step method), the promoter is initially separated from the antisense region by an intervening Kan^R^ element which is then subsequently deleted by site-specific excision between a pair of flanking FRT sites (81). The two integration events are designated as P*_tac_*::AS_b2459(Out) (in strain GJ20010/pHYD5701) and P*_tac_*::AS_b2459(In) (in strain GJ20013/pHYD5701), respectively.

Identical results were obtained following promoter integration by either method (that is expected to trigger antisense transcription in head-on conflict against replisome movement initiated from *oriC*), whereby the strain was now rendered RecA-dependent for viability [Fig. 3, panels i and ii, respectively, for (Out) and (In) integrants]; that is, only blue colonies were recovered in the blue-white assay described above. These findings indicate that P*_tac_*-directed antisense expression at the AS_b2459 locus leads to a compromise of genome integrity that necessitates RecA-mediated recombinational repair for survival.

**Figure 3:**
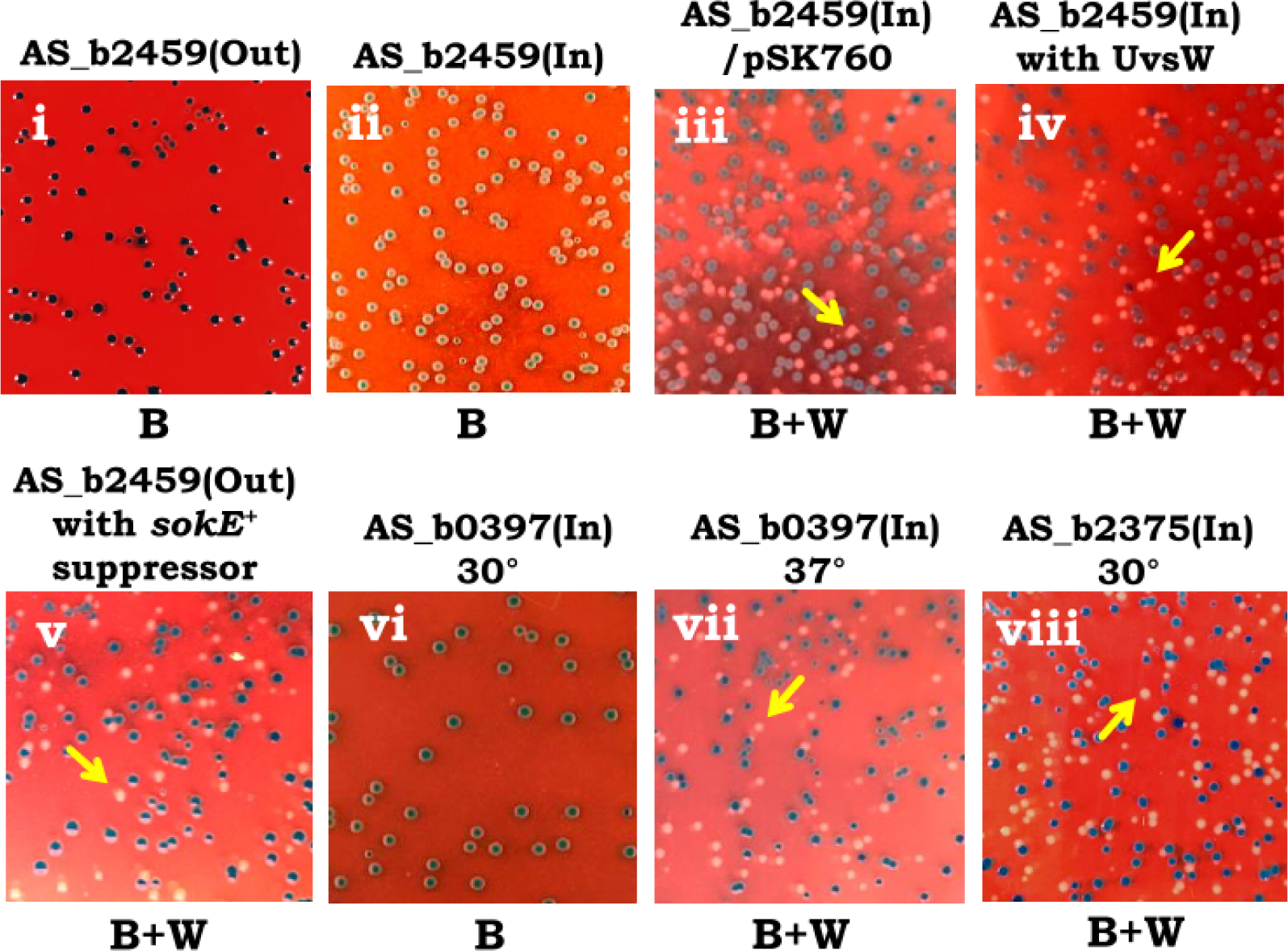
Blue-white assay of XL1-Blue derivatives with *recA^+^ lacZ^+^* shelter plasmid pHYD5701 to determine RecA-dependence or –independence for viability of chromosomal P*_tac_* integrants. Strains carrying chromosomally recombineered P*_tac_* at the loci as indicated were plated at suitable dilution on LB-Xgal and incubated for 24 h; representative images are shown withblue (B) or mixture of blue and white (B+W) colonies for the different derivatives, as marked beneath each panel. Examples of white colonies (signifying RecA-independence for viability) are marked by yellow arrows. The plates for panels vi and viii were incubated at 30°. The derivative for panel iii was plated on Amp-supplemented medium. Strains employed for the different panels werepHYD5701-carrying derivatives of: i, GJ20010; ii, GJ20013; iii, GJ20013/pSK760 (multicopy-*rnhA^+^*); iv, GJ20028; v, GJ20029; vi and vii, GJ20017; and viii, GJ20015.

Survival in absence of RecA (that is, growth of white colonies on Xgal medium) was restored for the P*_tac_*::AS_b2459(In) derivative (GJ20013) upon heterologous overexpression of RNase HI from a multicopy ColE1-derived plasmid pSK760 (Fig. 3, panel iii). Furthermore, in this situation, the *rnhA^+^*-bearing plasmid could no longer be cured from the *recA1* derivative even upon introduction of a second plasmid (pET28b, Kan^R^) of the same incompatibility group; thus, none of 40 Kan^R^ colonies tested had lost the Amp^R^-bearing *rnhA^+^* plasmid after introduction of the second Kan^R^ plasmid. On the other hand, when a similar experiment was performed as control in the isogenic *recA^+^* derivative (that is, on a blue colony), 32 of 40 Kan^R^ transformants tested had become Amp^S^, indicating that the *rnhA^+^* plasmid was being lost at high frequency as expected under these conditions. As with RNase HI overexpression, ectopic expression of the R-loop UvsW helicase (29, 43, 102) was also able to restore RecA-independent viability in the P*_tac_*::AS_b2459(In) integrant (Fig. 3, panel iv).

In a Δ*rnhA* derivative of XL1-Blue *recA^+^* (GJ20007/pHYD5701), attempts to introduce the P*_tac_*::AS_b2459(Out) event by Kan_R_ transduction were not successful. Likewise, in the method for obtaining P*_tac_*::AS_b2459(In) into this strain, the second step of site-specific excision of Kan^R^ (from strain GJ20022, which would result in P*_tac_* driving antisense expression at the locus) could not be achieved because of an inability to obtain transformants with plasmid pCP20 (that mediates the excision) into the strain. These results suggest that P*_tac_*-driven expression of AS_b2459 is lethal in a Δ*rnhA recA^+^* strain.

Taken together, these observations indicate that promoter integration at the chromosomal antisense locus AS_b2459 is correlated with generation of R-loops, and that the subsequent outcome is determined by cellular RNase HI status: with multicopy-*rnhA^+^*, the R-loops are removed without much ensuing pathology; in *rnhA^+^*, genome integrity is compromised most likely as a consequence of transcription-replication conflict at the site of R-loop formation (16, 48–50, 103–105), necessitating a requirement of recombinational repair for cell viability; and in Δ*rnhA*, there is persistent R-loop formation at the site that serves to permanently block replisome progression (48).

During the process of routine maintenance of cultures of these recombineered strains, we found that the synthetic lethality phenotype with *recA1* was often lost, suggestive of the accumulation of suppressor mutations. One such suppressor was designated GJ20029 (Fig. 3, panel v), and its molecular characterization (as a revertant of *sokE*::IS*150* to *sokE^+^*) is described in a later section.

### R-loops from engineered antisense transcription at chromosomal locus AS_b0397

For the second antisense region AS_b0397 (which confers lethality when expressed from plasmid pTrc99a only in a Δ*rnhA* background), the P*_tac_* promoter sequence was recombineered by the two-step method described above (to generate strain GJ20017/pHYD5701). The integration event is designated as P*_tac_*::AS_b0397(In). In this case, antisense transcription is expected to be co-directional with *oriC*-initiated replisome progression.

In the blue-white screening assay with *recA_+_* borne on the shelter plasmid, the P*_tac_* promoter integrant designed to express antisense region AS_b0397 failed to yield white colonies at 30° (Fig. 3, panel vi), indicating that this event also affects genome integrity as to be synthetic lethal with *recA1*. White colonies were obtained, however, at 37° (Fig. 3, panel vii). Previous studies have shown that R-loop toxicity is exacerbated at lower temperatures (42, 63).

### Absence of toxicity from engineered antisense transcription at control chromosomal locus AS_b2375

The P*_tac_* promoter was also recombineered for antisense transcription at the control chromosomal locus AS_b2375 by the two-step method described above (to generate strain GJ20015/pHYD5701), and the integration event is designated as P*_tac_*::AS_b2375(In). As determined by the blue-white screening assay with *recA^+^* on the shelter plasmid, both *recA^+^* and *recA1* derivatives were viable with 1 mM IPTG at 30° and 37°, confirming the absence of significant genome perturbation under these conditions (Fig. 3, data are shown for 30° in panel viii).

### Escape replication of defective prophages upon chromosomal overexpression of the R-loop prone antisense loci

We undertook WGS of cells from IPTG-supplemented cultures grown to exponential phase of the strains (all carrying the *recA^+^* plasmid) with P*_tac_* integrated upstream of the different antisense loci: P*_tac_*::AS_b2459(Out) and P*_tac_*::AS_b2459(In); P*_tac_*::AS_b0397(In) (two replicate cultures); and, as control, P*_tac_*::AS_b2375(In) (two cultures grown at, respectively, 37° and 30°). Cultures of the integrant at AS_b0397 were grown at 30° (since its phenotype of RecA-dependent viability is exhibited only at this temperature), whereas the pair of integrants at AS_b2459 was grown at 37°. DNA copy number for each of the regions across the entire chromosome was then determined in cultures of the test strains (with P*_tac_* integrated upstream of the R-loop prone antisense loci) relative to that in the control cultures of the P*_tac_*::AS_b2375(In) integrant grown at the cognate temperature.

The most interesting result from these experiments was that, in all the four test cultures that were RecA-dependent for viability, there was (in comparison with the control cultures) an enrichment in DNA copy number at one or both of a pair of genomic regions which precisely correspond to the positions of two defective (also called as cryptic or “grounded”) lambdoid prophages CP4-6 and DLP12 (106, 107); the two prophages are located at, respectively, 0.28 Mbp and 0.57 Mbp in the genome, and their positions are marked by red and green ovals in the different panels of Figure 4. Thus, for the P*_tac_* integrants at locus AS_b2459, the copy number of DLP12 prophage was elevated in one culture (In) (Fig. 4A) and that of CP4-6 in the other (Out) (Fig. 4B). For the integrant at locus AS_b0397, copy numbers of both CP4-6 and DLP12 were elevated in one replicate culture (Fig. 4C) whereas that of CP4-6 alone was elevated in the other (Fig. 4D). Careful examination of the plots also showed that the copy number of other defective prophages such as e14, CP4-44 AND CP4-57 were also modestly elevated in the culture of P*_tac_*::AS_b2459(Out) that was RecA-dependent for viability (Fig. 4B).

**Figure 4:**
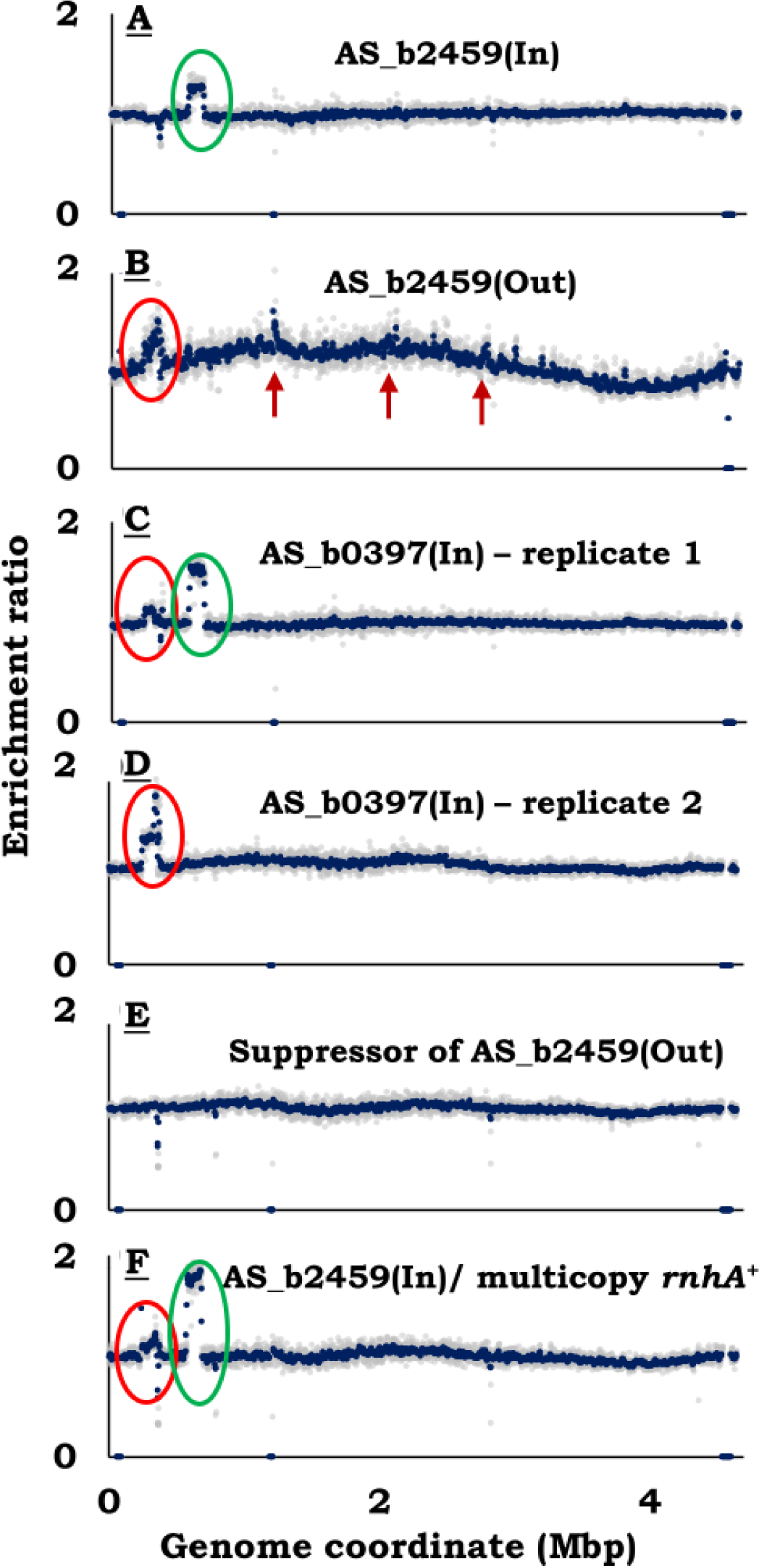
Chromosomal DNA copy number analysis in P*_tac_* integrant strains by WGS. (**A** to **F**) In each of the panels, DNA copy number distribution across the genome is shown for the indicated strain, as an enrichment ratio relative to that in the control strain. The control strain was GJ20015 [P*_tac_*::AS_b2375(In)] grown at either 37° (panels A, B, E and F) or 30° (panels C, D). Genomic positions of defective prophages CP4-6 and DLP12 are marked by red and green ovals, respectively. Arrows in Fig. 4B denote the positions of defective prophages (from left): e14, CP4-44 and CP4-57. Strains for the different panels were: A, GJ20013; B, GJ20010; C and D, GJ20017; E, GJ20029; and F, GJ20013/pSK760 (multicopy-*rnhA^+^*).

The copy number increases described above are indicative of escape replication (also called onion-skin replication) of the defective prophages in a subpopulation of cells in the cultures (108, 109). The mechanisms for regulation of their replication are not fully understood, and there is evidence that several including CP4-6 and DLP12 are controlled by the Rho factor (35, 37). Positive feedback regulatory loops, as in phage λ replication (110), may also be involved to give rise to bistable or all-or-none behaviour, which may account for differential amplification at the two prophage loci in replicate cultures. Strain background features may also play a role, as we show below that in the MG1655 background it is another defective prophage CPS-53 that exhibits pronounced escape replication upon Rho inhibition.

We suggest that it is the elevated synthesis of antisense transcripts from the identified R-loop prone loci, caused either by Rho deficiency or by P*_tac_* integration in a *rho^+^* strain, that triggers (in *trans*) escape replication of defective prophages as well as (in *cis*) R-loop associated arrest of RNAP leading to transcription-replication conflict (40, 44). A possible model that may link these two phenomena is discussed below.

### WGS analysis of suppressors of RecA-dependent viability

We also performed WGS on two suppressor derivatives of the P*_tac_*::AS_b2459 integrants in each of which RecA was no longer essential for viability. The first was a spontaneous mutant of P*_tac_*::AS_b2459(Out) that had arisen in the course of routine maintenance of the strain, and the WGS analysis indicated that the elevation in DNA copy number at the CP4-6 prophage locus that was observed in the parent had been reversed in this suppressor (Fig. 4E).

From the WGS reads of the parent and of this suppressor strain, we also performed de novo genome assembly and determined novel junction sequences by the method of Durrant et al (88). This analysis showed that whereas the parent had an IS*150* insertion in the *sokE* gene, the suppressor was *sokE^+^*; we then performed P1 transduction experiments to demonstrate that it is the *sokE* locus that is responsible for the difference in RecA dependence for viability between parent and the suppressor strain, and these results are described in the Supplementary Text. Analysis of WGS data of the other sequenced strains in this study showed that all of them were like the parent in harbouring an IS*150* insertion in *sokE*. Replacement by P1 transduction of *sokE*::IS*150* by *sokE^+^* in the other P*_tac_* integrants that were RecA-dependent for viability [P*_tac_*::AS_b2459(In) and P*_tac_*::AS_b0397(In)] also resulted in restoration of RecA-independent viability in them (described in the Supplementary Text).

The *E. coli* chromosome bears five *hok-sok* gene pairs (suffixed *A* to *E*) that encode orthogonal toxin-antitoxin systems which are all putatively inactivated in different ways (111, 112). The product of each of the *sok* genes (including *sokE*) is an antitoxin RNA of around 60 nucleotides. Several laboratory *E. coli* strains such as MG1655 and BW25113 are *sokE^+^*, but the gene is disrupted by IS*150* insertion (exactly as in our parental version of XL1-Blue) in more than 400 sequenced natural *E. coli* isolates, as also in some strains of *Shigella flexneri* [(113–115); see Supp. Table S4]; *sokA* of MG1655 is also disrupted by IS*150* insertion (112). Although the mechanism by which *sokE^+^* acts as a suppressor was not further investigated in this study, our results do suggest that P*_tac_*-directed chromosomal antisense expression, RecA-dependent viability, and prophage escape replication are indeed correlated with one another.

The second suppressor derivative (which was viable in absence of RecA) that was subjected to WGS was that with RNase HI overexpressed from a plasmid in the P*_tac_*::AS_b2459(In) integrant. In this case, the DNA copy number at both the CP4-6 and DLP12 prophage loci was elevated (Fig. 4F), although the parent itself had shown copy number increase only for DLP12; there was also the expected increase in read numbers mapping to the *rnhA* locus because of this gene’s additional presence on a multicopy plasmid (not discernible in Fig. 4F, in view of the small size of the locus). As discussed below, these findings indicate that although escape replication of the defective prophages may be correlated with elevated transcription of the R-loop prone antisense loci, it is likely not the cause for RecA-dependence of viability in the integrant strain.

In summary, the WGS analysis in combination with the phenotype data has shown that P*_tac_*-directed expression of R-loop forming antisense loci on the chromosome is associated with both escape replication of defective prophages and RecA dependence for strain viability, and that RNase HI overexpression apparently mitigates the latter but not the former.

### Increased prevalence of R-loops following Rho inhibition in C. crescentus and X. oryzae

To test whether the models developed for Rho’s role in R-loop avoidance in *E. coli* are extendable to other bacteria, we examined reactivity of total nucleic acid preparations from cultures of *C. crescentus* and *X. oryzae* pv. *oryzae* to the R-loop specific monoclonal antibody S9.6. Rho is essential for viability in both bacteria (116, 117). For *E. coli,* it has earlier been shown (29) that S9.6 reactivity is markedly increased in cultures grown in presence of a sublethal concentration of BCM, an inhibitor of Rho function in many bacterial species (4).

For both *C. crescentus* and *X. oryzae*, we first confirmed their growth sensitivity to BCM at, respectively, 100 and 30 μg/ml. Nucleic acid preparations were then obtained from the cultures grown with sublethal BCM (25 and 15 μg/ml, respectively), as also for *E. coli* MG1655 as control (with BCM at 25 μg/ml). These preparations exhibited strong signals with S9.6 antibody compared to those from the control cultures without BCM (Fig. 5A top panel, compare columns 1 and 3 from left). In each case, the S9.6 signal was considerably reduced or attenuated upon prior treatment of the preparations with RNase H in vitro (Fig. 5A top panel, compare columns 3 and 4 from left), indicating that the signal was indeed from RNA-DNA hybrids in the preparations. We conclude that R-loop abundance is enhanced upon Rho inhibition in both bacteria, just as is the case in *E. coli*.

**Figure 5:**
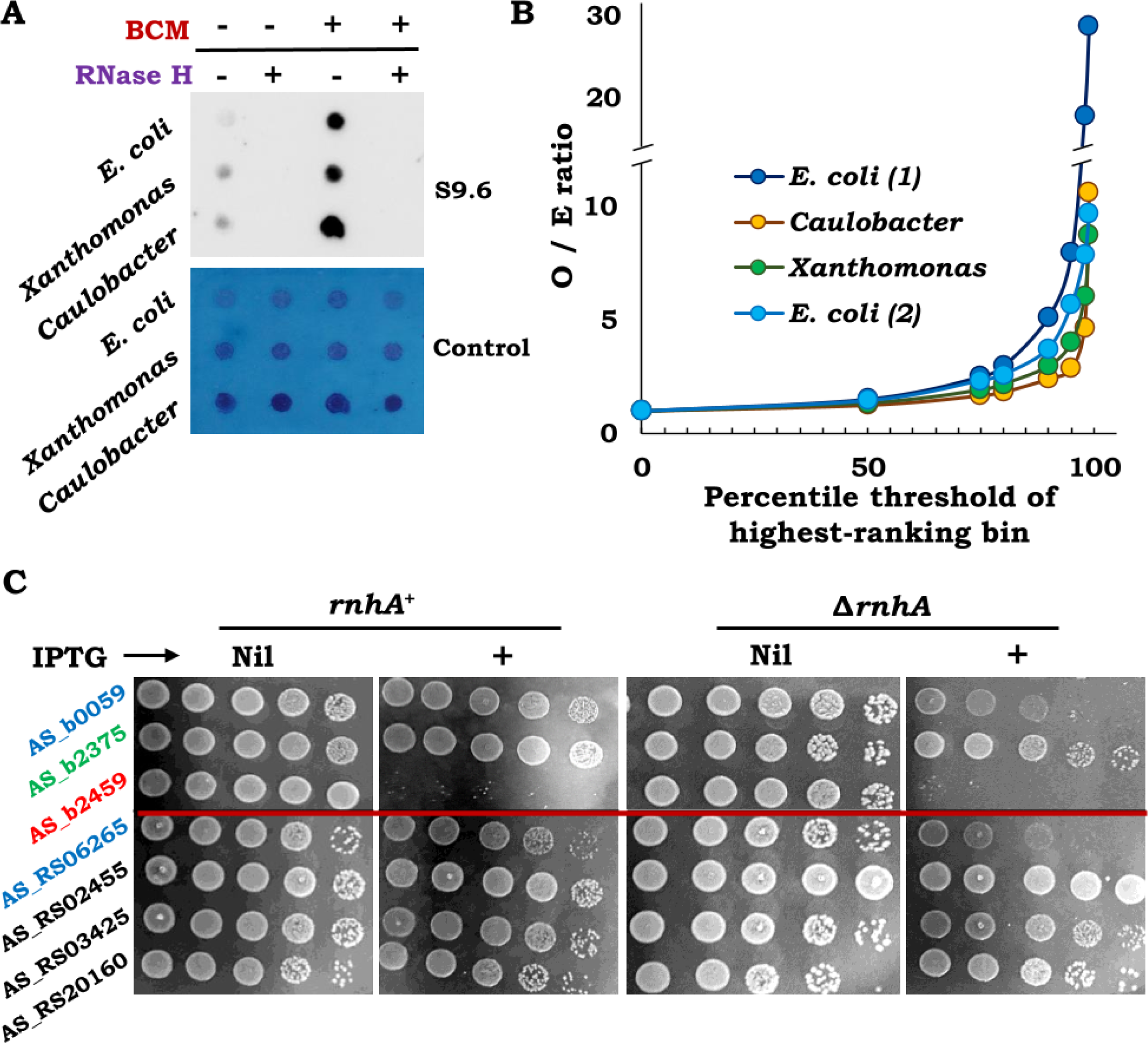
Putative R-looped antisense regions in *X. oryzae* and *C. crescentus*. (**A**) Top panel, Immunoblot with S9.6 monoclonal antibody of total nucleic acid preparations from indicated wild-type strains grown without or with sublethal BCM; treatment with RNase H, where indicated, was done in vitro. Bottom panel, methylene blue-stained image of the blotted membrane (to serve as loading control). (**B**) Concordance for antisense loci with high bisulphite reactivity between pairs of cultures, as indicated in the key. Each of the curves is a plot of the observed-to-expected (O/E) ratio for the highest-ranking bins of the different quantile ranges, as explained in the text. For the curves marked *Caulobacter*, *Xanthomonas*, and *E. coli (1)*, the comparisons were each between the pair of cultures (grown without and with BCM in this study) of the respective bacteria; for the curve marked *E. coli (2)*, the comparison was between *E. coli* cultures of this study and of an earlier study (43), both without BCM. (**C**) Dilution-spotting assays of pHYD5701 (*recA^+^*)-carrying derivatives of XL1-Blue (*rnhA^+^*) or GJ20007 (Δ*rnhA*), after transformation with pTrc99a derivatives carrying different cloned *X. oryzae* antisense regions; each of the numbers shown is prefixed with “AS_BXO1_” (as given in Supp. Table S2). Colour codes are as described in legend to Fig. 1. Two pTrc99a derivatives with cloned *E. coli* antisense regions are included as controls.

### Whole-genome bisulphite mapping for ss-DNA distribution in C. crescentus and X. oryzae

As mentioned above, we had previously employed a whole-genome bisulphite mapping strategy in *E. coli* to identify antisense loci with high propensity for R-loop formation (43), and we undertook a similar approach in this study with *C. crescentus* and *X. oryzae.* Non-denatured total nucleic acid preparations from cultures of the two bacteria (along with *E. coli* as control) grown without or with sublethal BCM were treated with sodium bisulphite, which leads to C-to-T conversions in ss-DNA. The preparations were then subjected to WGS in order to infer the distribution of ss-DNA regions in the genome. In this set of experiments, we could confirm the finding from an earlier report (36) that DNA copy number of a 10-kb region corresponding to the defective prophage CPS-53 (or KplE1), at the 2.47-Mbp location in the genome, is markedly increased in the Rho-inhibited *E. coli* culture, that is, with BCM supplementation (Supp. Table S5, sheet 1). These data are again indicative of prophage escape replication, similar to that described above for the RecA-dependent P*_tac_* integrant strains.

In addition to ss-DNA regions at R-loops which are exposed on the non-template strand behind the transcription elongation complex, ss-DNA is also expected to be present on the lagging-strand template during replisome progression. Accordingly, the ranking of bisulphite-reactive regions across the genome was done as in our previous study (43), with genes or ORFs as units, across four separate categories based on combinations of the following: whether the strand is the non-template strand for sense or for antisense transcription, and whether it is the leading-or lagging-strand template during replication. The data for the individual ORFs of *E. coli*, *X. oryzae* pv. *oryzae* and *C. crescentus* are presented in Supplementary Table S5 (sheets 1, 2 and 3, respectively).

As had been done in our earlier study (3), we then sought to identify the putative sites of antisense R-loop occurrence in the genome as those ORFs that rank high in bisulphite reactivity percentile for the non-template strand of antisense transcription. For each of the three pairs of cultures (of *E. coli*, *X. oryzae* and *C. crescentus*, without and with BCM), the *R^2^* correlation coefficient across the full range of antisense bisulphite reactivity percentiles was relatively modest at 0.51, 0.26 and 0.15, respectively. On the other hand, the correlations became progressively more robust at the higher percentile ranges, as was shown by the following analysis.

For each culture (say, *E. coli* without BCM), the antisense bisulphite reactivity data were grouped into different quantile ranges such as percentile, quintile, decile, dodecile, quartile, and so on. From each such range, the highest-ranking bin was chosen for comparison with its equivalent from the data of the partner culture (in this case, *E. coli* with BCM). Examples of the highest-ranking bins chosen were >99 for percentile, >95 for quintile, >90 for decile, and so on; and for each comparison, the number of identical ORFs observed in both sets, as a proportion of that expected by chance, was determined.

As shown in Figure 5B, the observed-to-expected ratio for identical ORFs within each of the pairs, computed as above, increases sharply with increasing threshold of the highest-ranking bin for all three bacteria. The same pattern was noted when the data from this study was compared with that from our earlier study (43) for *E. coli* cultures without BCM supplementation (Fig. 5B, curve marked *E. coli* (*2*); *R^2^* = 0.33 for the comparison between the full range of values).

This analysis suggests that, for all three bacteria, the subset of ORFs with high antisense bisulphite reactivity is highly reproducible. Based on the other findings that we have obtained in this and in earlier work (29, 43), we believe that at this high percentile range, R-loops are the main and reproducible source of antisense bisulphite reactivity. The data also lend support to our earlier proposal (43) that the order of ranking of R-loop prevalence across different antisense regions is not significantly different between cultures that are proficient or deficient for Rho-dependent transcription termination.

### Evidence for R-loop toxicity upon expression of X. oryzae antisense region in *E. coli*

Our work above had shown that around one-third (7/22) of the tested *E. coli* antisense regions, selected on the basis of high bisulphite reactivity percentile, confer toxicity associated with R-loop formation upon cloning and expression in plasmid pTrc99a. We undertook similar experiments with four selected antisense regions each from *X. oryzae* and *C. crescentus* that exhibited high bisulphite reactivity percentiles.

Of the total of eight antisense regions that were tested from the two organisms after cloning in plasmid pTrc99a (listed in Supp. Table S2), one (from *X. oryzae*, AS_BXO1_RS06265) conferred an IPTG-dependent decrease of viability in the Δ*rnhA E. coli* mutant but not in the *rnhA^+^* parent (both carried the *recA^+^* plasmid pHYD5701); the other seven clones were viable with IPTG in both strains (Fig. 5C, data are shown only for derivatives with cloned *X. oryzae* antisense regions). These findings lend support to the notion that at least some of the antisense regions exhibiting high bisulphite reactivity from these organisms are prone to R-loop formation upon their adventitious expression in *E. coli*.

## Discussion

Synthesis of the vast majority of antisense RNAs in *E. coli* is kept under tight check by Rho-and NusG-mediated transcription termination (29, 30). We have previously presented evidence for the occurrence of two categories of novel antisense transcripts in Rho-deficient cells, namely those that form and do not form R-loops, respectively. The loci AS_b2459 and AS_b0397 encode examples of the first category whereas locus AS_b2375 encodes an example of the second. In the present work, we have examined the effects of engineered overexpression of these two categories of Rho-curtailed antisense RNAs (in Rho-proficient strains) to show that those of the first category generate toxic R-loops both on a plasmid and on the chromosome.

In eukaryotes as well, the Sen1/senataxin factor has been proposed to function much like Rho in bacteria in mediating both transcription termination and prevention of R-loop occurrence from nascent transcripts (118–122). More generally, it is being increasingly recognized that various cotranscriptional processes are important for gene regulation in both prokaryotes and eukaryotes (123).

### R-loop generation on plasmids from *E. coli* antisense RNAs

Of the 22 candidate R-loop forming antisense regions that were selected based on the data from whole-genome studies of bisulphite reactivity, we obtained evidence of R-loop toxicity conferred by seven of them (including AS_b2459 and AS_b0397) upon their cloning and expression from plasmid pTrc99a. (None of the three control antisense regions of the second category, including AS_b2375, conferred toxicity in these experiments.) These represent the first examples in which occurrence of R-loops from endogenous DNA regions cloned on a plasmid has been demonstrated in wild-type *E. coli*; in previous studies, R-loop formation on plasmids has been shown either in topoisomerase I-deficient *E. coli* with some endogenous DNA regions (64, 100), or in wild-type *E. coli* with heterologous DNA stretches from other organisms (92, 93).

We also note that the antisense regions that confer R-loop toxicity upon ectopic expression from plasmid pTrc99a were rather sharply demarcated, and that in several instances even adjacent antisense regions (which otherwise fulfilled the candidate criteria) did not confer similar toxicity. Two alternative explanations, not mutually exclusive, for these findings are (i) that initiation of R-loop formation (as opposed to R-loop extension) occurs at specific sequences (124), and that plasmid toxicity occurs when sites of such R-loop initiation are cloned and expressed; or (ii) that the end-point of cell lethality used in the assay represents a high threshold for R-loop detection, and that some antisense regions that also generate R-loops fail to meet this threshold level.

Our studies have shown that two other bacteria *X. oryzae* pv. *oryzae* and *C. crescentus* are sensitive to the Rho inhibitor antibiotic BCM, and that R-loop prevalence in them is increased in presence of sublethal BCM just as is the case in *E. coli* (29). For these organisms as well, we believe that at least some of the identified antisense regions with high-percentile bisulphite reactivity would represent sites of R-loop formation, which was indeed confirmed for one such locus after its cloning and expression in plasmid pTrc99a.

### Escape replication of defective prophages and RecA-dependent cell viability upon P*_tac_*-directed chromosomal expression of R-loop prone antisense regions

P*_tac_*-directed chromosomal overexpression of the R-loop prone (test) antisense loci AS_b2459 and AS_b0397 (but not of the control antisense locus AS_b2375) was associated both with RecA-dependence for cell viability as well as with escape replication of defective prophages. Upon RNase HI overexpression in the strain with antisense transcription of the AS_b2459 chromosomal locus, cell viability in absence of RecA was restored but prophage escape replication did persist. We propose the following model to explain these findings.

We suggest that the fundamental difference between the two categories of antisense loci whose transcription is provoked in Rho-deficient cells is that only one of them is very susceptible to translation-uncoupled RNAP backtracking and arrest (40, 58, 59, 125); it is in this category (that includes AS_b2459 and AS_0397) then that R-loops are generated consequent to arrest of the elongating RNAP, leading in *cis* to serious transcription-replication conflict and the need for RecA-mediated recombinational repair to maintain cell viability.

We further propose that escape replication of defective prophages is an event that is triggered in *trans* following the backtracking and arrest of RNAP on this category of antisense loci. The phenomenon is an accompaniment to, but independent of, R-loop formation, and the signal linking escape replication to RNAP backtracking remains to be identified. Some possibilities are that the anti-backtracking factors GreA/GreB and/or participants of the DksA-ppGpp axis play a role in such signal generation (40, 126, 127).

According to our model, when RNase HI is overexpressed in cells with P*_tac_*-directed expression of AS_b2459, the R-loops associated with the arrested transcription elongation complex are removed so that transcription-replication conflict is mitigated. On the other hand, the backtracked RNAP without the R-loop persists (which does not pose a barrier to replication since it is easily dislodged during replisome progression) and hence the postulated signal for prophage escape replication continues to be generated.

### Disruption of genome integrity through site-specific R-loop formation on the chromosome

Co-transcriptional R-loop formation has previously been implicated as an obstacle to DNA replication and as a threat to genome integrity in both bacteria and eukaryotic cells (44, 48, 49, 51, 97, 128–130). It would appear that both co-directional and head-on transcription-replication conflict associated with R-loop formation can affect genome integrity (40, 48, 49, 104, 105, 131), which is supported in the present study as well. Of the two recombineered antisense loci, one is expected to generate head-on (AS_b2459) and the other co-directional (AS_b0397) transcription-replication conflict.

The ability to generate site-specific DNA damage on the bacterial chromosome has been useful in experimental studies on the mechanisms both of damage generation and of its repair. Previously described methods for this purpose have included placement of I-SceI sites (86, 132), ectopic *tus* sites (75), a palindrome sequence for cleavage by SbcCD (76), or sites for binding of LacI or TetR protein arrays (73, 74). We suggest that the recombineered antisense R-looped loci generated in this work may also serve as suitable tools for use in such studies.

## Data availability

All genome sequence data described in this work are available for full public access at https://www.ncbi.nlm.nih.gov/bioproject/PRJNA1101904.

## Supplementary data

Supplementary data are provided in a PDF file “Supplementary data”, along with two additional Excel files as Tables S4 and S5, respectively.

## Acknowledgements

We thank Anjana Badrinarayanan, Subhadeep Chatterjee and R Harinarayanan for strains; and Papri Basak, Rachna Chaba, Manjula Ekka and Abhijit Sardesai for advice and discussions.

## Supplementary Text

### Cloning of chromosomal antisense regions into plasmid pTrc99a

PCR primers for amplifying the chromosomal antisense regions from *E. coli*, *X. oryzae* pv. *oryzae*, and *C. crescentus* carried restriction sites for directional cloning into pTrc99a. The PCR amplicons were treated with the appropriate pair of restriction enzymes and ligated with pTrc99a DNA that had been similarly digested before transformation into strain XL1-Blue. Details for cloning each of the regions are given in Supplementary table S2, and the primer sequences used for PCR in Supplementary Table S3.

### Integration by recombineering of P*_tac_* at different chromosomal locations

Construction of the pKD13 derivatives pHYD7527 and pHYD7528, carrying the cloned P*_tac_* sequence in two orientations, has been described in the main text. PCR amplicons for integration of P*_tac_* by recombineering into the different chromosomal loci were prepared as follows (primer sequences for PCR are given in Supp. Table S3): (i) P*_tac_*::AS_b2459(Out), primer pair JGAH073-JGAH074 on plasmid template pHYD7527; (ii) P*_tac_*::AS_b2459(In), primer pair JGAH075-JGAH076 on plasmid template pHYD7528; (iii) P*_tac_*::AS_b0397(In), primer pair JGAH112-JGAH113 on plasmid template pHYD7528; (iv) P*_tac_*::AS_b2375(In), primer pair JGAH110-JGAH111 on plasmid template pHYD7528; (v) P*_tac_*::*lacZ*(Out), primer pair JGAH105-JGAH106 on plasmid template pHYD7527; and (vi) P*_tac_*::*lacZ*(In), primer pair JGAH103-JGAH104 on plasmid template pHYD7528.

For each of the PCR amplicons from (i) to (iv) above, recombineering was achieved by selection for Kan^R^ following transfection into strain HR421 (that carries the mini-/\-Tet prophage), and the integrant allele was then transferred by P1 transduction into XL1-Blue carrying the *recA^+^* plasmid pHYD5701. For the PCR amplicons from (v)-(vi) above, transfections were done into MG1655/pKD46 (1) with selection for Kan^R^, and the integrant alleles were then transferred by P1 transduction into MG1655; in both cases, the recombineering events lead to replacement by P*_tac_* of the native *lac* promoterupstream of the *lacZYA* operon of MG1655, to yield integrants in the (Out) and (In) orientations, respectively.

### Measurement of magnitude of leaky expression from P*_tac_* integrated in the chromosome

To determine the strength of expression from chromosomally integrated P*_tac_* without and with IPTG supplementation, we measured the specific activity of β-galactosidase [in Miller units (2)] in MG1655 and its P*_tac_* integrant derivatives at the *lacZYA* locus (pairs of values given for each strain are without and with IPTG supplementation, respectively): MG1655 (with native *lac* promoter), 18 and 1050; GJ20023 [P*_tac_*::Kan^R^-*lacZYA*(In), where Kan^R^ element is located between P*_tac_* and Kan^R^], 3 and 90; GJ20025 [P*_tac_*::FRT-*lacZYA*(In), in which the Kan^R^ element between P*_tac_* and *lacZYA* has been removed by site-specific excision], 219 and 1964; and GJ20024 [P*_tac_*::*lacZYA*(Out), where P*_tac_* reads away from Kan^R^ directly into *lacZYA*], 258 and 1996.

### Confirmation by P1 transductions of *sokE* as a suppressor of RecA-dependent viability in chromosomal P*_tac_* integrants

As described in the main text, a comparison between the WGS data of the strain GJ20010 with P*_tac_*::AS_b2459(Out) and of its suppressor derivative GJ20029 (that had become RecA-independent for viability) had indicated that the former carries an IS*150* insertion in *sokE* whereas the latter is *sokE^+^*. To determine whether this difference in the *sokE* locus between the two strains is responsible for suppression, we performed transductions to Kan^R^ with phage P1 lysates prepared on three strains from the Keio knockout collection (Δ*entE*, Δ*cstA*, and Δ*ilvD*) into the Kan^S^ derivative of the P*_tac_*::AS_b2459(Out) parental strain (GJ20011). The Keio collection strains are derivatives of BW25113 which is *sokE^+^*, and the loci *entE* and *cstA* are situated, respectively, 19 kb and 23 kb away from *sokE*; the *ilvD* locus is distant from *sokE* and was chosen as control in these experiments.

The Kan^R^ transductants obtained were tested in the blue-white assay for determining whether they retained the parental phenotype of RecA-dependence for viability (that is, yielding only blue colonies with the *recA^+^*-bearing shelter plasmid pHYD5701) or had become RecA-independent (yielding blue and white colonies). For the control transduction with Δ*ilvD*::Kan, all transductants tested remained RecA-dependent for viability (thereby establishing that the recipient culture was free of spontaneous suppressor mutants). On the other hand with Δ*entE*::Kan and Δ*cstA*::Kan, 4 of 12 and 5 of 12 transductants, respectively, had become RecA-independent. Since there are no other DNA sequence differences in this region between the donor and recient strains in the P1 transductions, we conclude that it is the *sokE* locus that determines the RecA requirement for viability in the P*_tac_*::AS_b2459(Out) strain.

We also performed similar transduction experiments to Kan_R_ with the three P1 lysates into the P*_tac_*::AS_b2459(In) and P*_tac_*::AS_b0397(In) strains (GJ20013 and GJ20017, respectively) to establish that *sokE_+_* suppresses the RecA requirement for viability in both of them. The following were the proportions of Kan^R^ transductants that yielded white colonies in these crosses with the P1 lysates prepared on Δ*ilvD*, Δ*entE* and Δ*cstA*, respectively: for P*_tac_*::AS_b2459(In), 0/3, 4/12, and 4/12; and for P*_tac_*::AS_b0397(In), 0/3, 6/12, and 3/12.

---

**Table S1:** List of *E. coli* strains.

| Strain No <sup>a</sup> | Genotype <sup>b</sup> |
| --- | --- |
| MG1655 | *E. coli* K-12 wild-type |
| XL1-Blue | *recA1 endA1 gyrA96 thi-1 hsdR17 supE44 relA1 lac [F' proAB lacI<sup>q</sup>ZΔM15 Tn10] sokE::IS150* |
| HR421 | MG1655 Δ*araBAD mal::lacI<sup>q</sup>* mini-λ-Tet |
| GJ20001 | XL1-Blue Δ*pcnB*::Kan |
| GJ20007 | XL1-Blue Δ*rnhA*::Kan |
| GJ20008 | XL1-Blue Δ*rnhA*::FRT |
| GJ20009 | GJ20008 Δ*pcnB*::Kan |
| GJ20010 <sup>c</sup> | XL1-Blue P<sub>tac</sub>::AS_b2459(Out)(Kan) |
| GJ20011 <sup>c</sup> | XL1-Blue P<sub>tac</sub>::AS_b2459(Out)(FRT) |
| GJ20013 <sup>c</sup> | XL1-Blue P<sub>tac</sub>::AS_b2459(In)(FRT) |
| GJ20015 | XL1-Blue P<sub>tac</sub>::AS_b2375(In)(FRT) |
| GJ20017 <sup>c</sup> | XL1-Blue P<sub>tac</sub>::AS_b0397(In)(FRT) |
| GJ20022 | GJ20008 P<sub>tac</sub>::AS_b2459(In)(Kan) |
| GJ20023 | XL1-Blue P<sub>tac</sub>::*lacZYA*(In)(Kan) |
| GJ20024 | XL1-Blue P<sub>tac</sub>::*lacZYA*(Out)(Kan) |
| GJ20025 | XL1-Blue P<sub>tac</sub>::*lacZYA*(In)(FRT) |
| GJ20028 | GJ20013 *att λ*::P<sub>ara</sub>-UvsW (Amp) |
| GJ20029 | *sokE*<sup>+</sup> spontaneous suppressor of GJ20010 |

<sup>a</sup> Previously described strains include MG1655 (3) and XL-1 Blue (4). Strain HR421 was kindly provided by R Harinarayanan.
<sup>b</sup> For the strains constructed in this study, the knockout alleles Δ*pcnB*::Kan and Δ*rnhA*::Kan were sourced from the Keio collection (3), and the *att λ*::P<sub>ara</sub>-UvsW (Amp) allele has been described earlier (5). Alleles from which the Kan<sup>R</sup> marker has been excised by pCP20-mediated recombination are shown with the suffix '::FRT'.
<sup>c</sup> Each of the indicated strains was maintained in the stock collection as a derivative with a *recA*<sup>+</sup>-carrying shelter plasmid pHYD5701.

**Table S2:**
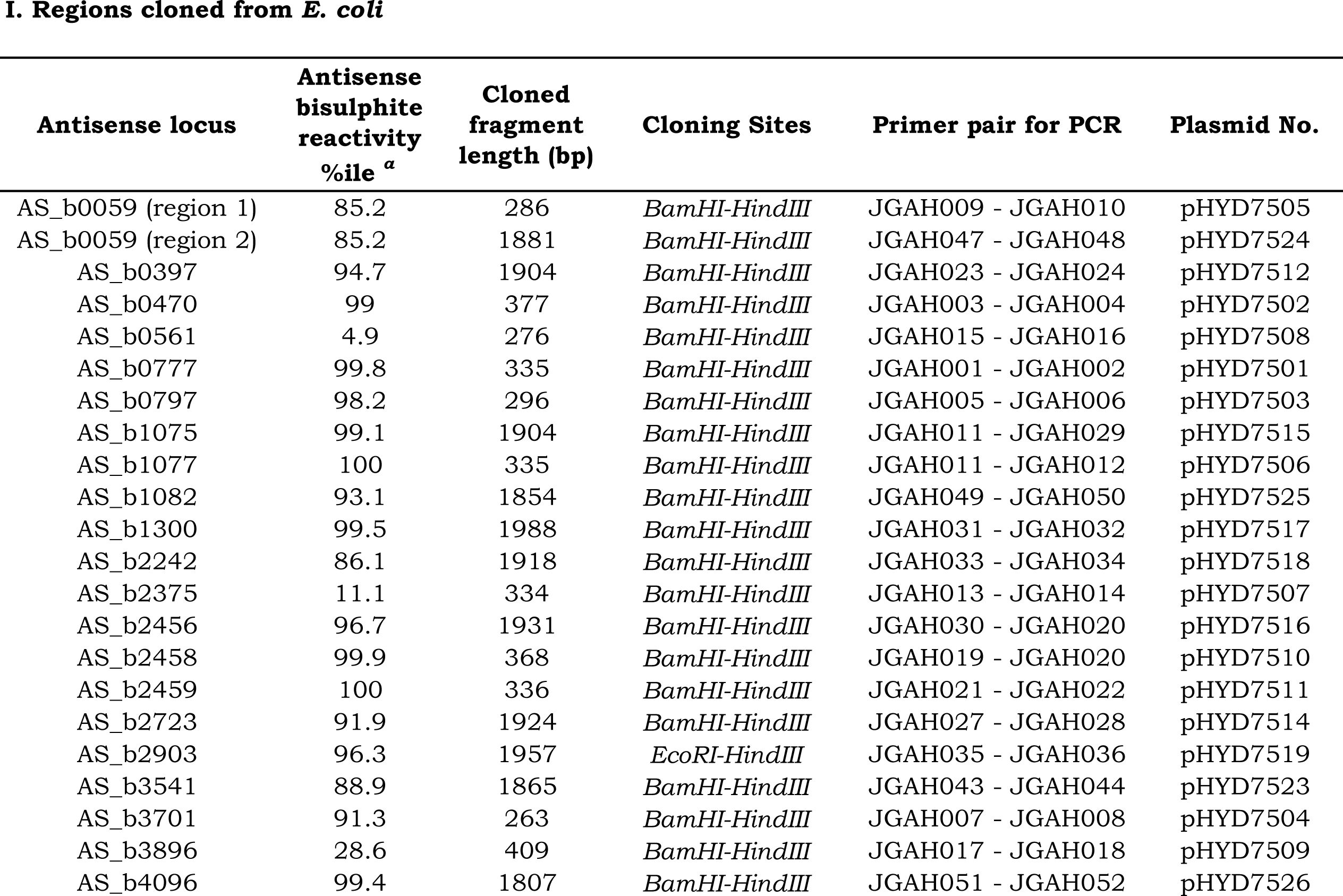
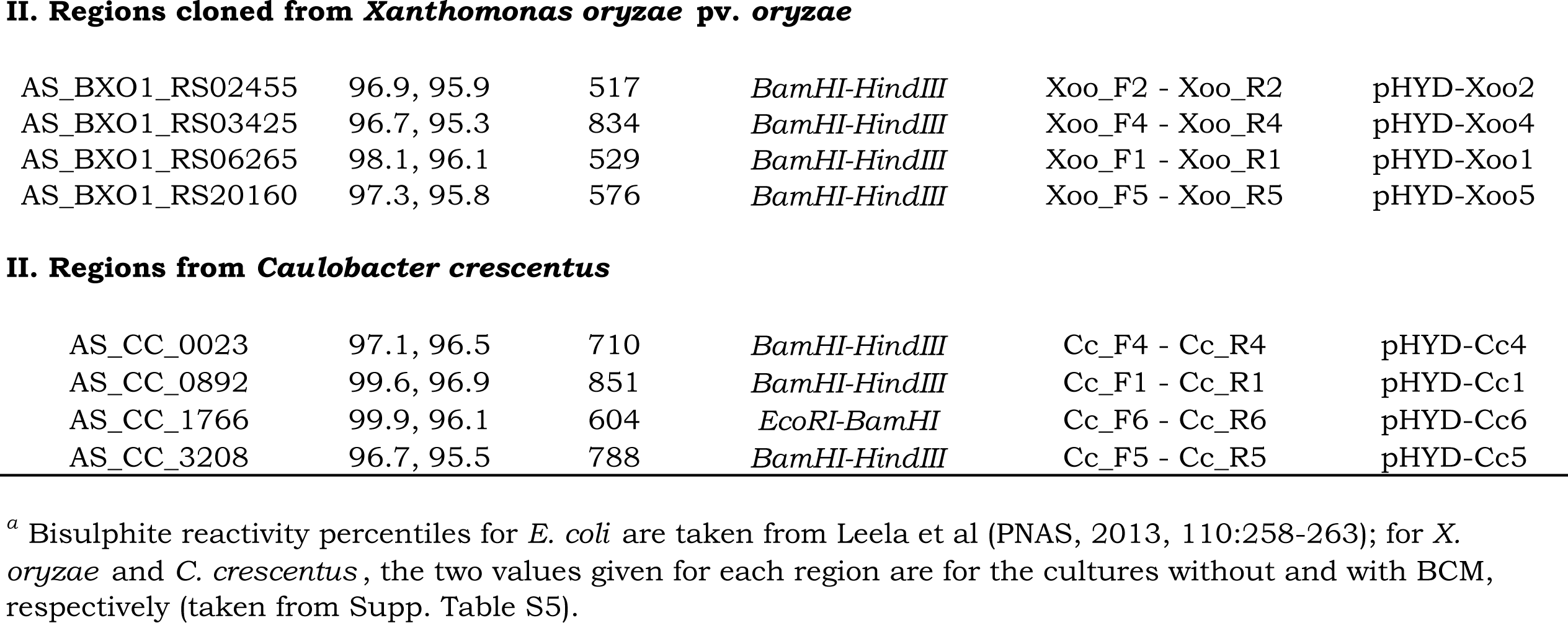
List of pTrc99A-derived plasmids.

Regions cloned from *E. coli*
| **Antisense locus** | **Antisense**<br>**bisulphite**<br>**reactivity**<br>**%ile <sup>a</sup>** | **Cloned**<br>**fragment**<br>**length (bp)** | **Cloning Sites** | **Primer pair for PCR** | **Plasmid No.** |
| --- | --- | --- | --- | --- | --- |
| AS_b0059 (region 1) | 85.2 | 286 | *Bam*HI-*Hind*III | JGAH009 - JGAH010 | pHYD7505 |
| AS_b0059 (region 2) | 85.2 | 1881 | *Bam*HI-*Hind*III | JGAH047 - JGAH048 | pHYD7524 |
| AS_b0397 | 94.7 | 1904 | *Bam*HI-*Hind*III | JGAH023 - JGAH024 | pHYD7512 |
| AS_b0470 | 99 | 377 | *Bam*HI-*Hind*III | JGAH003 - JGAH004 | pHYD7502 |
| AS_b0561 | 4.9 | 276 | *Bam*HI-*Hind*III | JGAH015 - JGAH016 | pHYD7508 |
| AS_b0777 | 99.8 | 335 | *Bam*HI-*Hind*III | JGAH001 - JGAH002 | pHYD7501 |
| AS_b0797 | 98.2 | 296 | *Bam*HI-*Hind*III | JGAH005 - JGAH006 | pHYD7503 |
| AS_b1075 | 99.1 | 1904 | *Bam*HI-*Hind*III | JGAH011 - JGAH029 | pHYD7515 |
| AS_b1077 | 100 | 335 | *Bam*HI-*Hind*III | JGAH011 - JGAH012 | pHYD7506 |
| AS_b1082 | 93.1 | 1854 | *Bam*HI-*Hind*III | JGAH049 - JGAH050 | pHYD7525 |
| AS_b1300 | 99.5 | 1988 | *Bam*HI-*Hind*III | JGAH031 - JGAH032 | pHYD7517 |
| AS_b2242 | 86.1 | 1918 | *Bam*HI-*Hind*III | JGAH033 - JGAH034 | pHYD7518 |
| AS_b2375 | 11.1 | 334 | *Bam*HI-*Hind*III | JGAH013 - JGAH014 | pHYD7507 |
| AS_b2456 | 96.7 | 1931 | *Bam*HI-*Hind*III | JGAH030 - JGAH020 | pHYD7516 |
| AS_b2458 | 99.9 | 368 | *Bam*HI-*Hind*III | JGAH019 - JGAH020 | pHYD7510 |
| AS_b2459 | 100 | 336 | *Bam*HI-*Hind*III | JGAH021 - JGAH022 | pHYD7511 |
| AS_b2723 | 91.9 | 1924 | *Bam*HI-*Hind*III | JGAH027 - JGAH028 | pHYD7514 |
| AS_b2903 | 96.3 | 1957 | *Eco*RI-*Hind*III | JGAH035 - JGAH036 | pHYD7519 |
| AS_b3541 | 88.9 | 1865 | *Bam*HI-*Hind*III | JGAH043 - JGAH044 | pHYD7523 |
| AS_b3701 | 91.3 | 263 | *Bam*HI-*Hind*III | JGAH007 - JGAH008 | pHYD7504 |
| AS_b3896 | 28.6 | 409 | *Bam*HI-*Hind*III | JGAH017 - JGAH018 | pHYD7509 |
| AS_b4096 | 99.4 | 1807 | *Bam*HI-*Hind*III | JGAH051 - JGAH052 | pHYD7526 |
| II. Regions cloned from *Xanthomonas oryzae* pv. *oryzae* |  |  |  |  |  |
| AS_BXO1_RS02455 | 96.9, 95.9 | 517 | *Bam*HI-*Hind*III | Xoo_F2 - Xoo_R2 | pHYD-Xoo2 |
| AS_BXO1_RS03425 | 96.7, 95.3 | 834 | *Bam*HI-*Hind*III | Xoo_F4 - Xoo_R4 | pHYD-Xoo4 |
| AS_BXO1_RS06265 | 98.1, 96.1 | 529 | *Bam*HI-*Hind*III | Xoo_F1 - Xoo_R1 | pHYD-Xoo1 |
| AS_BXO1_RS20160 | 97.3, 95.8 | 576 | *Bam*HI-*Hind*III | Xoo_F5 - Xoo_R5 | pHYD-Xoo5 |
| II. Regions from *Caulobacter crescentus* |  |  |  |  |  |
| AS_CC_0023 | 97.1, 96.5 | 710 | *Bam*HI-*Hind*III | Cc_F4 - Cc_R4 | pHYD-Cc4 |
| AS_CC_0892 | 99.6, 96.9 | 851 | *Bam*HI-*Hind*III | Cc_F1 - Cc_R1 | pHYD-Cc1 |
| AS_CC_1766 | 99.9, 96.1 | 604 | *Eco*RI-*Bam*HI | Cc_F6 - Cc_R6 | pHYD-Cc6 |
| AS_CC_3208 | 96.7, 95.5 | 788 | *Bam*HI-*Hind*III | Cc_F5 - Cc_R5 | pHYD-Cc5 |

<sup>a</sup> Bisulphite reactivity percentiles for *E. coli* are taken from Leela et al (PNAS, 2013, 110:258-263); for *X. oryzae* and *C. crescentus*, the two values given for each region are for the cultures without and with BCM, respectively (taken from Supp. Table S5).

**Table S3:**
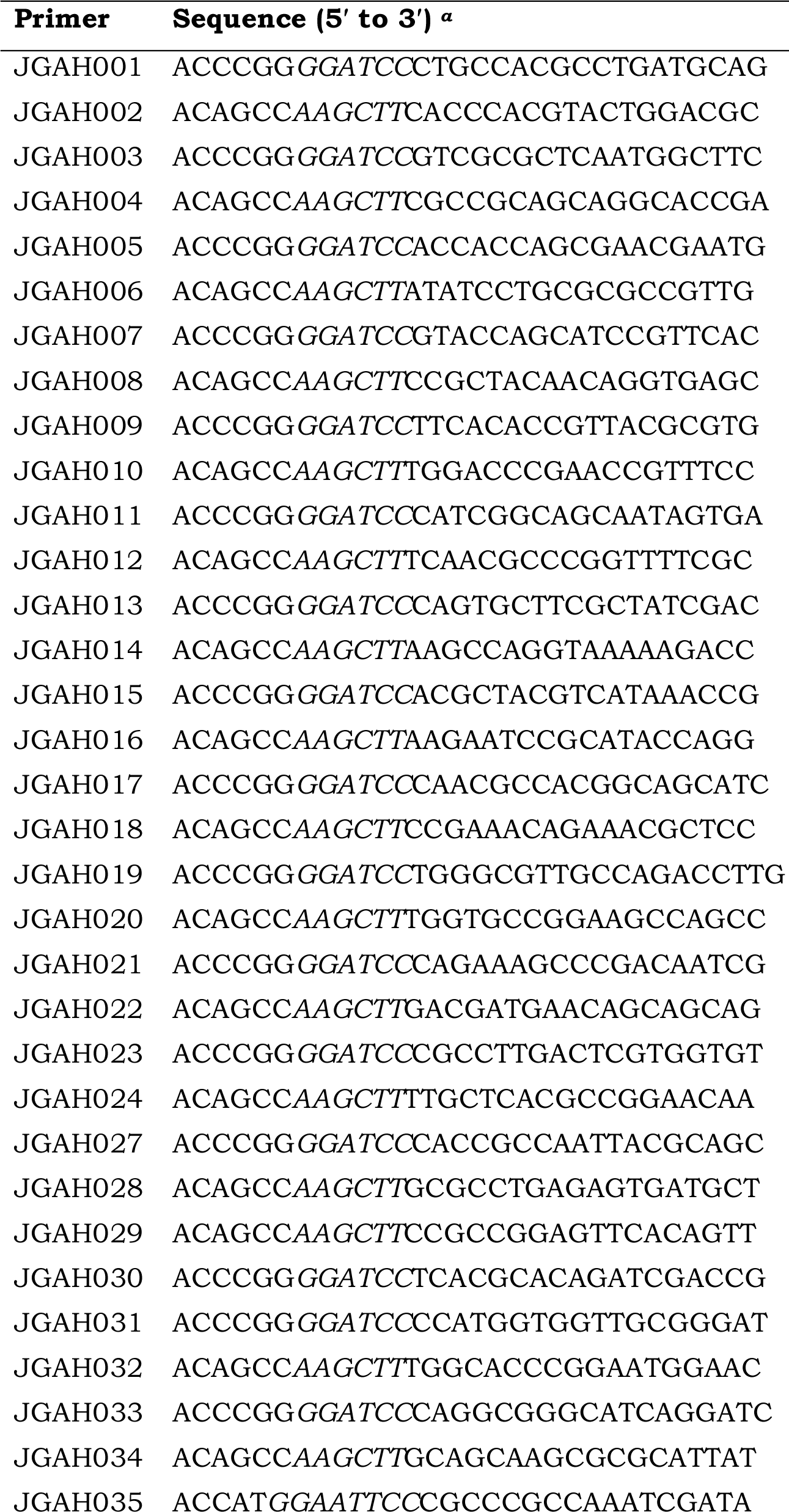
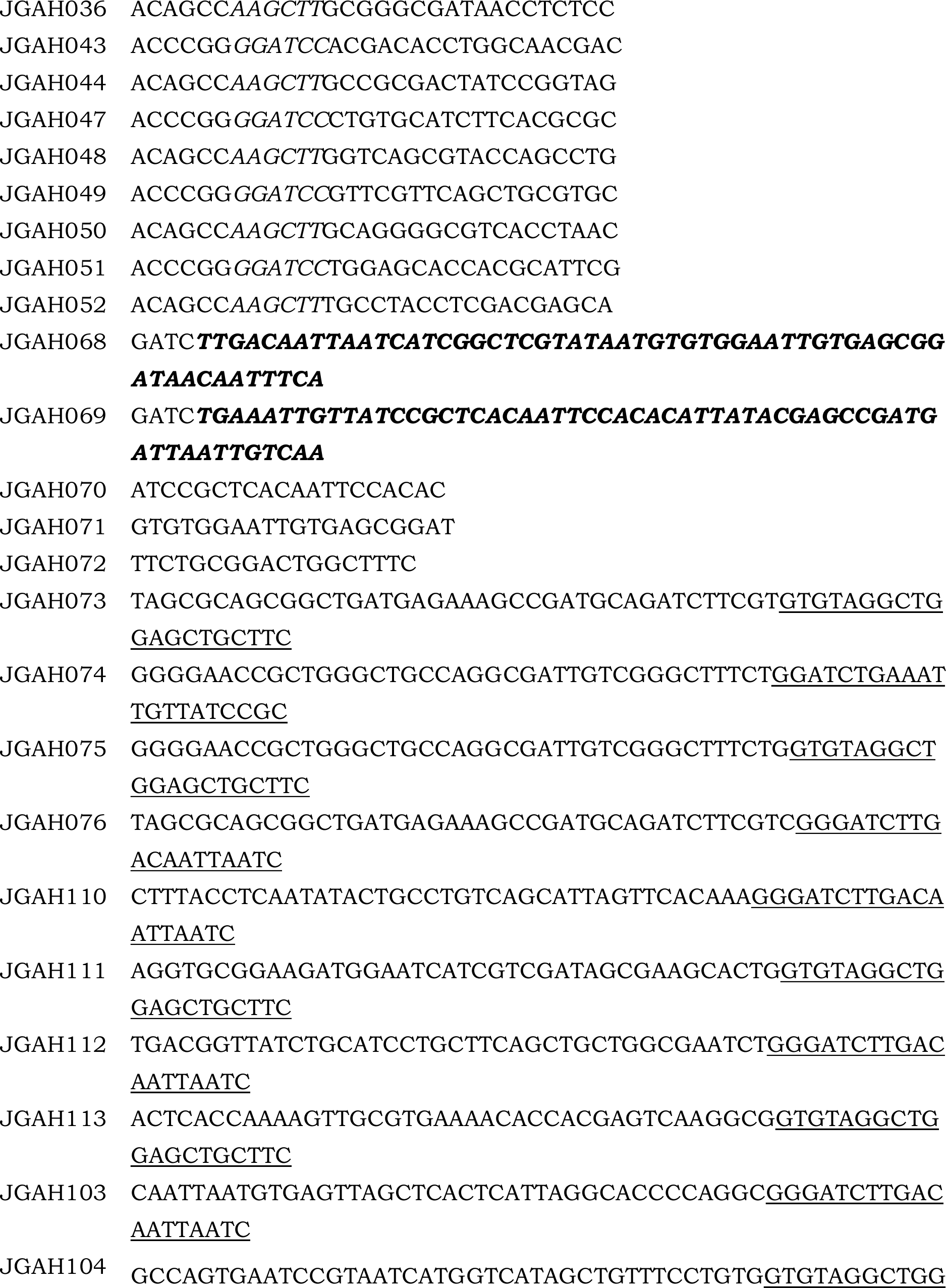
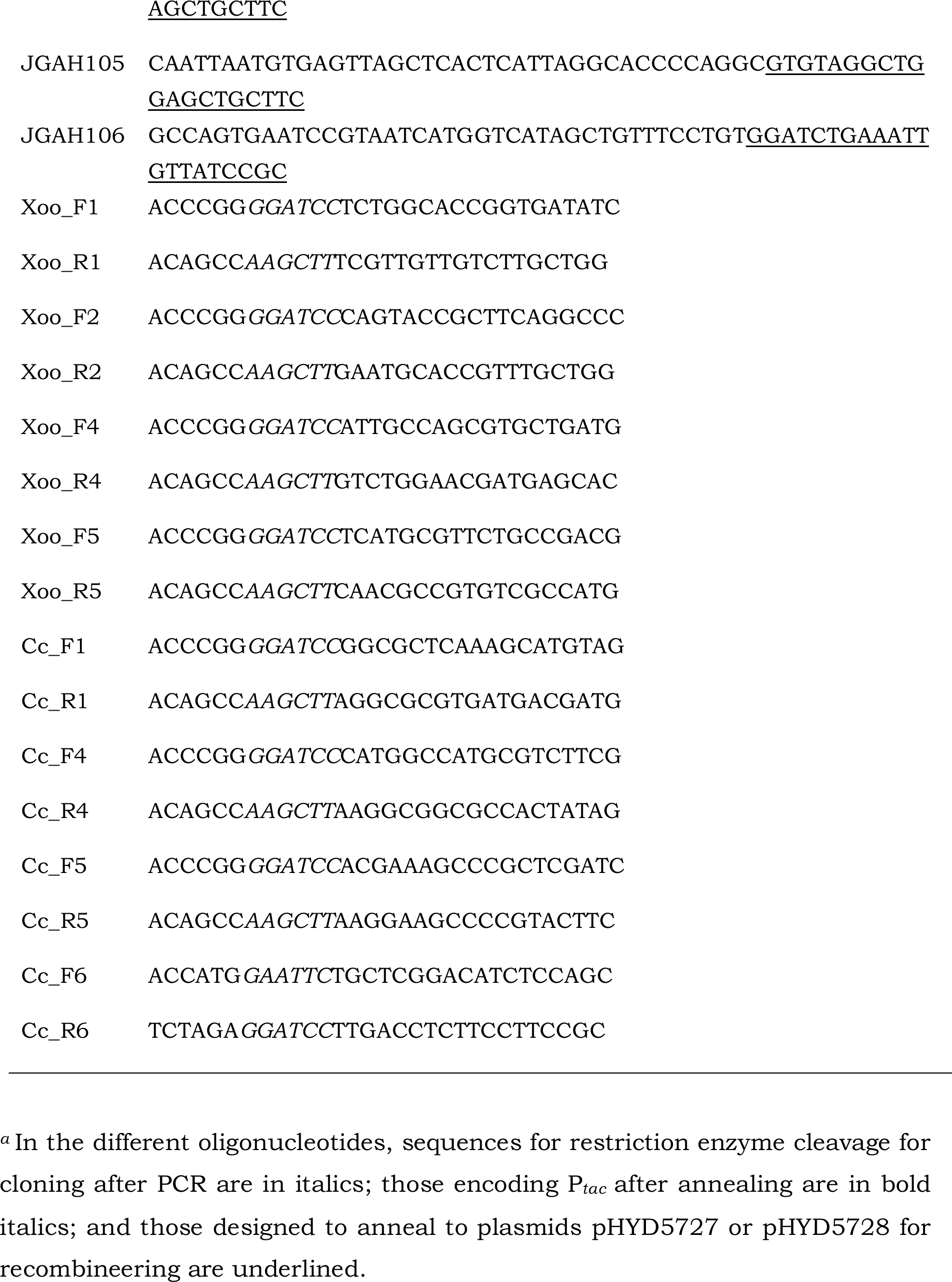
List of oligonucleotide primers.

**Supp. Fig. S1:**
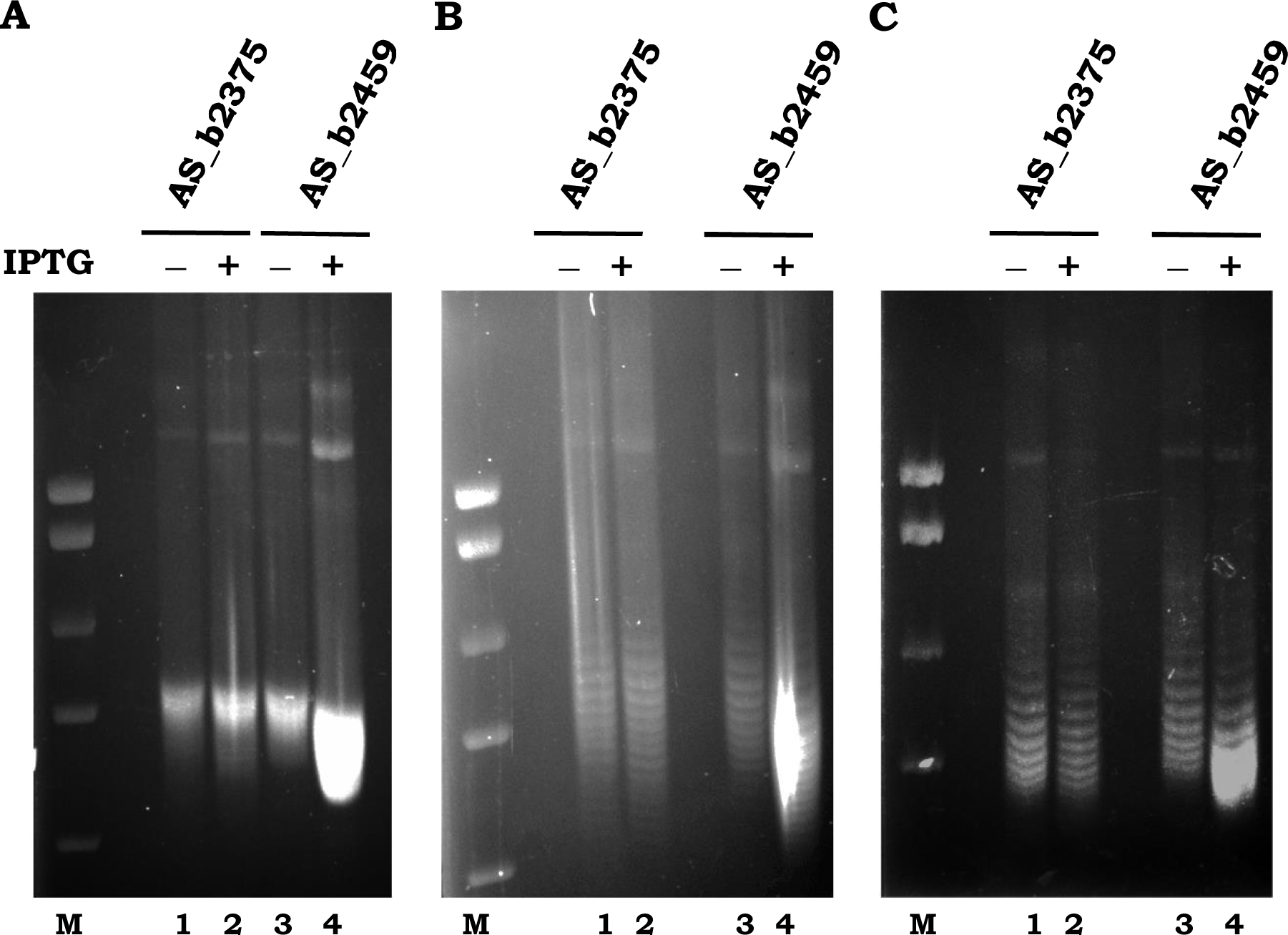
(**A** to **C**) Supercoiling status of pTrc99a derivatives with cloned antisense regions AS_b2497 (test) and AS_b2375 (control) in cultures without and with IPTG supplementation. Gel electrophoresis was on 1% agarose with chloroquine added at 4, 6 and 7 μg/ml, respectively. Notations are the same as those described in the legend to Fig. 2.

